# Internalizing Symptoms Associate with the Pace of Epigenetic Aging in Childhood

**DOI:** 10.1101/776526

**Authors:** Marieke S. Tollenaar, Roseriet Beijers, Elika Garg, T.T. Thao Nguyen, David T.S. Lin, Julia L. MacIsaac, Idan Shalev, Michael S. Kobor, Michael J. Meaney, Kieran J. O’Donnell, Carolina de Weerth

**Affiliations:** Institute of Psychology, Leiden University, the Netherlands; Department of Developmental Psychology, Behavioural Science Institute, Radboud University, the Netherlands; Ludmer Centre for Neuroinformatics and Mental Health, Douglas Hospital Research Centre, McGill University, QC, Canada; Sackler Program for Epigenetics and Neurobiology, McGill University, QC, Canada; Centre for Molecular Medicine and Therapeutics, Department of Medical Genetics, BC Children’s Hospital Research Institute, University of British Columbia, BC, Canada; Department of Biobehavioral Health, Pennsylvania State University, PA, USA; Canadian Institute for Advanced Research, Child and Brain Development Program, Canada; Singapore Institute for Clinical Sciences, Singapore

**Keywords:** Childhood, Internalizing Symptoms, Externalizing Symptoms, Epigenetic Age Acceleration, Biological Aging

## Abstract

Childhood psychiatric symptoms may be associated with advanced biological aging. This study examined whether epigenetic age acceleration (EAA) associates with internalizing and externalizing symptoms across childhood in a longitudinal cohort study. At age 6 buccal epithelial cells from 148 children (69 girls) were collected to survey genome-wide DNA methylation. EAA was estimated using the Horvath clock. Internalizing symptoms at ages 2.5 and 4 years significantly predicted higher EAA at age 6, which in turn was significantly associated with internalizing symptoms from ages 6 to 10 years. Similar trends for externalizing symptoms did not reach significance. These findings indicate advanced biological aging in relation to child mental health and may help better identify those at risk for lasting impairments associated with internalizing disorders.

---

Common mental health disorders, such as anxiety and depression, result in chronic human suffering and come at a large economic cost (Whiteford et al., 2013). As psychiatric symptoms often have their origins early in life (Reef, Diamantopoulou, van Meurs, Verhulst, & van der Ende, 2009), early indicators of childhood psychiatric symptoms are of high importance. Recent studies show that depressive symptoms in childhood and adolescence may be associated with advanced physical aging, such as early pubertal and adrenarcheal timing (Copeland, Worthman, Shanahan, Costello, & Angold, 2019; Ellis et al., 2019; Lewis et al., 2018), possibly due to early life adversity (Gur et al., 2019). Here we examine whether a novel indicator of advanced biological aging based on genomic DNA methylation, i.e., epigenetic age acceleration (EAA), is associated with childhood psychiatric symptoms.

Epigenetic age has recently emerged as a cross-tissue index of biological aging (Horvath & Raj, 2018). EAA has been associated with cancer, cardiovascular disesase and all-cause mortality, independent of chronological age (Chen et al., 2016; Marioni et al., 2015; Perna et al., 2016), although not in all studies (Kim et al., 2018; Murabito et al., 2018). EAA is furthermore predicted by chronic life stress (Gassen, Chrousos, Binder, & Zannas, 2017; Zannas et al., 2015), and has been associated with mental health problems in adult populations (Fries et al., 2017; Han et al., 2018; Wolf & Schnurr, 2016).

First studies indicate that EAA is also an indicator of advanced physical maturation in younger populations, as demonstrated by early pubertal development and higher cortisol output in adolescence (Binder et al., 2018; Davis et al., 2017; Suarez, Lahti, Czamara, Lahti-Pulkkinen, Girchenko, et al., 2018). A longitudinal study in a pediatric sample from birth to age 17 also suggests that EAA is associated with advanced maturation in certain physical domains, e.g. fat mass and height, although not in others (e.g. not with puberty) (Simpkin et al., 2017). Importantly, EAA is also associated with higher odds for psychiatric problems in adolescent girls (Suarez, Lahti, Czamara, Lahti-Pulkkinen, Girchenko, et al., 2018). However, associations between EAA and psychiatric symptoms in younger children remain unknown.

We generated EAA estimates from buccal cells at age 6 in a longitudinal study sample, and examined whether it associates with trajectories of internalizing and externalizing symptoms across childhood. We first examined whether early childhood symptoms predicted EAA at age 6, and secondly whether EAA at age 6 would predict symptom development from ages 6 to 10.

## Method

### Participants

Participants are children of a Dutch ongoing longitudinal study including 193 mothers and children (Beijers, Jansen, Riksen-Walraven, & de Weerth, 2010; Tollenaar, Beijers, Jansen, Riksen-Walraven, & De Weerth, 2011). At age 6 years, genomic DNA was extracted from buccal epithelial cells in 148 of these children (69 girls, 79 boys). Mothers gave informed consent, and the local Institutional Ethics Committee, which follows the Helsinki Declaration, approved the study protocols. See the Supplementary information for further details on the study sample.

### Psychiatric Symptoms

Maternal reports were used to index childhood internalizing and externalizing symptoms. At ages 2.5 and 4 years the CBCL 1.5-5 (Achenbach & Rescorla, 2000) was used and at ages 6, 7 and 10 the CBCL 4-18 (Achenbach, 2007). Internalizing and externalizing symptoms at ages 8 and 10 were also measured with the SDQ (Mieloo et al., 2014), to confirm if study outcomes are consistent across symptom checklists. Continuous symptom scale scores were used in the analyses, but CBCL t-scores are included in Table 1 for reference.

**Table 1.** Descriptive statistics for internalizing and externalizing symptoms from ages 2.5 to 10 years.

|  | CBCL<sup>a</sup><br>Age 2.5 | CBCL<sup>a</sup><br>Age 4 | CBCL<sup>b</sup><br>Age 6 | CBCL<sup>b</sup><br>Age 7 | CBCL<sup>b</sup><br>Age 10 | SDQ<br>Age 8 | SDQ<br>Age 10 |
| --- | --- | --- | --- | --- | --- | --- | --- |
| N | 145 | 139 | 143 | 143 | 133 | 145 | 137 |
| *Internalizing Symptoms* |  |  |  |  |  |  |  |
| Mean | 7.20 | 6.33 | 4.09 | 4.61 | 5.60 | 3.21 | 2.80 |
| SD | 4.78 | 4.96 | 3.95 | 4.44 | 5.47 | 2.80 | 2.66 |
| Range | 0-21 | 0-24 | 0-21 | 0-24 | 0-35 | 0-14 | 0-15 |
| Mean t-score<br>(range) | 48.0<br>(29-66) | 46.0<br>(29-69) | 48.3<br>(32-73) | 49.4<br>(32-75) | 50.3<br>(33-80) |  |  |
| *Externalizing Symptoms* |  |  |  |  |  |  |  |
| Mean | 9.68 | 11.05 | 6.55 | 6.43 | 6.31 | 4.83 | 4.02 |
| SD | 4.88 | 7.07 | 5.23 | 5.73 | 5.96 | 3.47 | 3.29 |
| Range | 0-21 | 0-33 | 0-26 | 0-29 | 0-30 | 0-17 | 0-16 |
| Mean t-score<br>(range) | 50.8<br>(28-71) | 47.7<br>(28-73) | 49.3<br>(32-72) | 48.6<br>(32-74) | 50.5<br>(33-74) |  |  |

*Notes:* CBCL =Child Behavior Checklist, SDQ =Strengths and Difficulties Questionnaire,<sup>a</sup> The CBCL1.5-5 was used, <sup>b</sup> The CBCL4-18 was used. t-scores represent normative gender-adjusted values of the CBCL for the respective age groups.

We also considered maternal anxiety and depressive symptoms at ages 2.5, 4, 6, 8 and 10 to ensure our results were not confounded by maternal mood at time of reporting child symptoms. Maternal symptoms of anxiety and depression were assessed using the State-Trait Anxiety Inventory (STAI) (Spielberger, 1983) and the Edinburgh Postnatal Depression Scale (EPDS) (Pop, Komproe, & van Son, 1992) respectively.

### Epigenetic Age Acceleration (EAA) and genetic covariates

Genome-wide DNA methylation was assessed with the Infinium EPIC array and genotyping was performed using the Infinium Global Screening Array (see Supplementary information for details). Epigenetic age was calculated with the script developed by Horvath (2013). EAA was estimated by the residuals that result from regressing epigenetic age on chronological age. A positive EAA indicates higher than expected epigenetic age, whereas a negative EAA indicates lower than expected epigenetic age.

To control for cell heterogeneity (Zheng et al., 2018), we used an approach informed by Smith et al. (2015) to estimate buccal epithelial cell content and blood cell contamination (CD34+ cells). Since these showed a very high correlation (*r* =−.93, *p* <.001) only buccal epithelial cell content was included as a covariate. We used a well-established Principal Component (PC) analysis-based approach to control for population stratification (Price et al., 2006). From this analysis the first 2 PCs were identified to best describe the population structure of our cohort and were included as covariates. We also considered potential genetic risk factors for EAA based on studies in brain and blood (Lu et al., 2016, 2018). However, none of these genetic risk markers contributed to EAA (see Supplementary information) and were hence not included in further analyses.

### Statistical analyses

IBM SPSS 24 was used for all analyses with α at .05. Analyses that were separately performed for internalizing and externalizing symptoms had α corrected to .025. To account for the non-normal distributed CBCL and SDQ data, Spearman rank correlations were used in the descriptive analyses, and log-transformed values in the main analyses.

For the prediction of EAA the following covariates were examined: child sex, buccal cell content, 850K technical factors (batch number, array plate, array position), and population stratification (PC 1 and PC2). For the prediction of CBCL scores over time covariates included: child sex, maternal educational level at birth, mothers’ own anxiety and depression ratings per time point, and population stratification (PC 1 and PC2). All covariates were at once included in a first step in the regression and mixed models. Only significant covariates (*p* <.05) were kept in the main analyses to increase power by reducing the number of predictors in relation to the sample size.

Linear regression analyses were performed with internalizing and externalizing symptoms at ages 2.5 and 4 as predictors of EAA at age 6. Covariates were examined as described above. To examine whether increases in symptoms at age 4 compared to age 2.5 would associate with higher levels of EAA, scores at 2.5 and 4 were included together in a final regression model. Increases in *R*^2^ (∆*R*^2^) were used as indicators of model fit and reflected additionally variance explained per model.

Linear mixed model analyses were performed with EAA at age 6 as a predictor of internalizing and externalizing symptoms at ages 6, 7, and 10 assessed with the CBCL. Mixed models allow for the inclusion of time varying components, and for individual missing values between time points. The intercept was included as a random factor. In addition to the covariates, linear and curvilinear time components were examined in the first step to assess development of symptoms from age 6 to 10. Then EAA was added as a main effect, and subsequently in interaction with time to examine whether EAA would also predict an increase in symptom development over time. Lastly, internalizing and externalizing symptoms at ages 2.5 and 4 were included to examine whether EAA could predict symptoms from age 6 to 10 beyond early childhood symptoms alone. Changes in random intercept variance were used as an estimate of additionally explained variance between individuals. As confirmatory analyses, similar mixed model analyses were performed for internalizing and externalizing symptoms assessed with the SDQ at ages 8 and 10 as outcome measure.

## Results

The average chronological age of the participants at the time of DNA sampling was 6.09 years (*SD* =0.14, range: 5.87 – 6.85 years). The average epigenetic age was 4.64 years (*SD* =.80, range: 3.07 – 6.93 years), which was significantly lower than chronological age, *t*(147) =21.8, *p*<.001, and not correlated to it (*r* =.077, *p* =.36). Measures of EAA ranged from −1.58 to 2.21 years, with an average of 0 (*SD* =.80), and correlated very strongly with epigenetic age (*r* =.997, *p* <.001). Boys and girls did not differ in EAA, *t*(146)=−.30, *p* =.77.

Table 1 shows internalizing and externalizing symptoms from ages 2.5 to 10. One participant, who developed an autism spectrum disorder by age 10, had high internalizing and externalizing symptoms (>3 SD from the mean) on 3 occasions and was removed from subsequent analyses. No sex differences were found on the CBCL scales at any age (all *p*s >.07). At age 10 only, girls exhibited higher levels of SDQ-reported internalizing problems than boys (*F*[1, 135] =4.03, *p* =.04). Symptom scores over time were positively correlated (see Table 2), and internalizing and externalizing symptoms were positively correlated at all time points (see Supplementary TableS1). Table 2 also shows positive correlations between EAA at age 6 and psychiatric symptoms over time, indicating both cross-sectional and longitudinal positive associations, mainly for internalizing symptoms. Bivariate associations between internalizing and externalizing symptoms scores over time, EAA and all covariates can be found in Supplementary TableS2 and TableS3.

**Table 2.** Spearman rank correlations between CBCL and SDQ scales over time and EAA, for internalizing and externalizing symptoms separately.

| *Internalizing symptoms* |  |  |  |  |  |  |  |  |
| --- | --- | --- | --- | --- | --- | --- | --- | --- |
| Measure | Age | CBCL<br>2.5 | CBCL<br>4 | CBCL<br>6 | CBCL<br>7 | CBCL<br>10 | SDQ<br>8 | SDQ<br>10 |
| CBCL | 2.5 | 1 |  |  |  |  |  |  |
| CBCL | 4 | .58*** | 1 |  |  |  |  |  |
| CBCL | 6 | .42*** | .43*** | 1 |  |  |  |  |
| CBCL | 7 | .32*** | .40*** | .64*** | 1 |  |  |  |
| CBCL | 10 | .47*** | .48*** | .52*** | .53*** | 1 |  |  |
| SDQ | 8 | .19* | .23** | .37*** | .54*** | .49*** | 1 |  |
| SDQ | 10 | .32*** | .33*** | .35*** | .42*** | .56*** | .55*** | 1 |
| **EAA** | **6** | **.22\*\*** | **.20\*** | **.25\*\*** | **.20\*** | **.18\*** | **.22\*\*** | **.09** |
| *Externalizing symptoms* |  |  |  |  |  |  |  |  |
| Measure | Age | CBCL<br>2.5 | CBCL<br>4 | CBCL<br>6 | CBCL<br>7 | CBCL<br>10 | SDQ<br>8 | SDQ<br>10 |
| CBCL | 2.5 | 1 |  |  |  |  |  |  |
| CBCL | 4 | .56*** | 1 |  |  |  |  |  |
| CBCL | 6 | .44*** | .51*** | 1 |  |  |  |  |
| CBCL | 7 | .38*** | .53*** | .72*** | 1 |  |  |  |
| CBCL | 10 | .37*** | .49*** | .66*** | .69*** | 1 |  |  |
| SDQ | 8 | .32*** | .44*** | .43*** | .61*** | .49*** | 1 |  |
| SDQ | 10 | .30*** | .41*** | .46*** | .57*** | .57*** | .78*** | 1 |
| **EAA** | **6** | **.14** | **.11** | **.14** | **.18\*** | **.08** | **.11** | **.21\*** |

Notes: CBCL = Child Behavior Checklist, SDQ = Strengths and Difficulties Questionnaire, EAA = Epigenetic Age Acceleration, Age in years, \* $p < .05$ , \*\* $p < .01$ , \*\*\* $p < .001$ .

### Predicting EAA at age 6 from childhood psychiatric symptoms

Linear regression analyses were performed to predict EAA at age 6. Covariates were first examined as predictors of EAA. Array position was the only significant covariate (*b* =−.30, *p* =.006) and subsequently included in the regression models.

Internalizing symptoms at age 2.5 significantly predicted EAA (*b* =.20, *p* =.015, ∆*R*^2^=.04). Internalizing symptoms at age 4 also significantly predicted EAA (*b* =.19, *p* =.024, ∆*R*^2^=.035). When symptoms at age 4 were added to model with symptoms at age 2.5, no significant additional variance was explained (∆*R*^2^=.003, *p* =.53). See Supplementary TableS4 for an overview of the regression models. To visualize the association between early childhood internalizing symptoms and EAA, Figure 1A shows a scatterplot of the association between standardized internalizing symptoms at age 2.5 and EAA at age 6.

**Figure 1.**
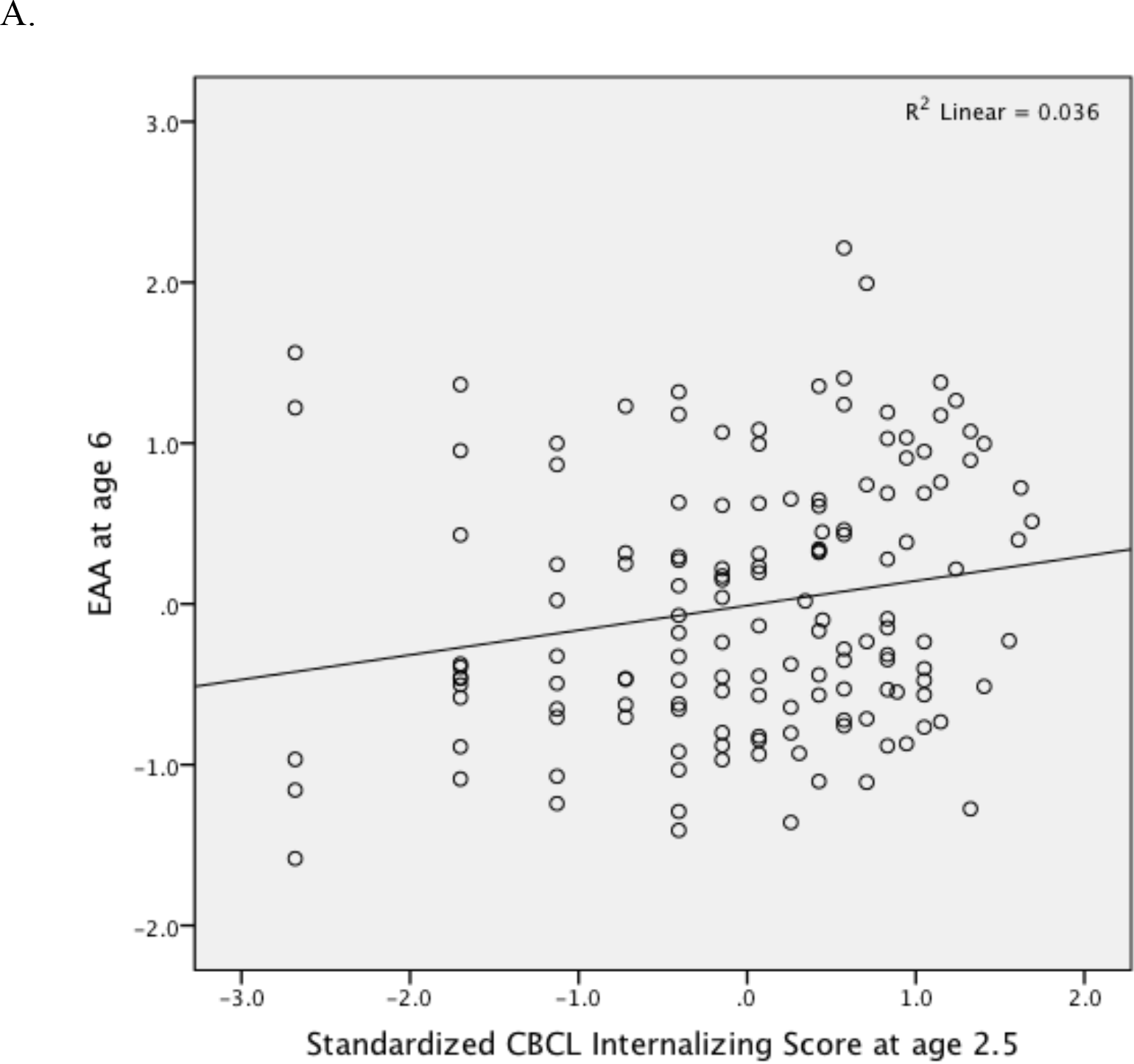
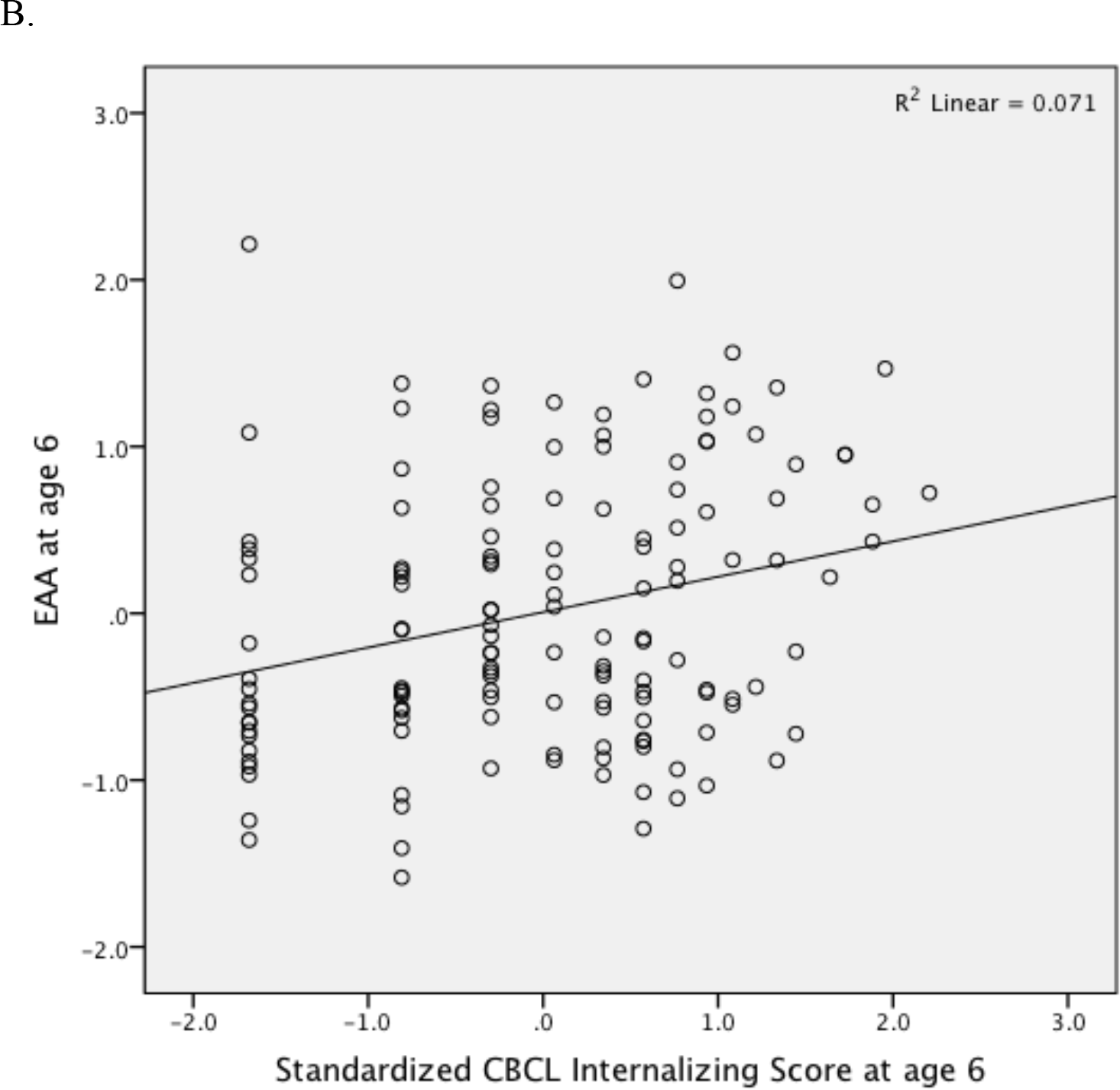
Associations between standardized internalizing scores at age 2.5 (A) and 6 (B) measured with the Child Behavior Checklist (CBCL) and Epigenetic Age Acceleration (EAA) at age 6.

Externalizing symptoms at age 2.5 did not significantly predict EAA (*b* =.12, *p* =.14, ∆*R*^2^ =.015), and neither did externalizing symptoms at age 4 (*b* =.12, *p* =.16, ∆*R*^2^=.015). When both externalizing symptoms at ages 2.5 and 4 were included, EAA was not significantly predicted by either (*p*s > .05, see Supplementary TableS5). Furthermore, when externalizing symptoms at age 2.5 were added to a model with internalizing symptoms at age 2.5, this did not account for additional variance in EAA (∆*R*^2^ < .001, *p* =.83; externalizing symptoms: *b* =.021, *p* =.83, internalizing symptoms: *b* =.18, *p* =.065).

### Predicting childhood psychiatric symptoms over time from EAA at age 6 Internalizing symptoms

Covariate analyses showed that time as a linear factor (beta =.048, SE =.014, *t* =3.54, *p* <.001), PC1 (beta =−.901, SE =.346, *t* =−2.60, *p* =.01)^2^ and maternal STAI scores (beta =.0096, SE =.0026, *t* =3.66, *p* <.001) significantly predicted CBCL internalizing symptoms from age 6 to 10, indicating an increase in internalizing symptoms over time, and positive associations between maternal anxiety problems and her report on child internalizing symptoms.

EAA was associated with significantly higher levels of internalizing symptoms from age 6 to 10 (beta =.096, SE =.028, *t* =3.42, *p* =.001), explaining 9.4% of variance between individuals. The interaction of EAA with time was not significant (*p* >.65, see Supplementary TableS6 for an overview of these mixed model analyses). We then examined whether EAA would predict internalizing symptoms from age 6 to 10 *in addition* to symptoms earlier in childhood. Symptoms at age 2.5 significantly predicted higher levels of symptoms at age 6 to 10 (beta =.125, SD =.022, *t* =5.61, *p* <.001). EAA remained a significant predictor in addition to symptoms at age 2.5 (beta =.074, SE =.027, *t* =2.77, *p* =.006), explaining an additional 6.7% of variance between individuals, indicating that EAA and early childhood symptoms are at least in part independent predictors of later childhood internalizing symptoms. When symptoms at both age 2.5 and 4 years were considered in a model together with EAA, EAA was associated with later child symptoms at trend level (beta =.050, SE =.027, *t* =1.88, *p* =.063), explaining an additional 3.1% of variance between individuals, see Supplementary TableS7.

In line with the CBCL data, EAA also showed a significant positive association with SDQ-assessed internalizing scores from age 8 to 10 after controlling for relevant covariates (beta =.065, SD =.026, *t* =2.46, *p* =.015), explaining 4.9% of variance between individuals. Again no interaction of EAA with time was found (beta =−.0047, SD =.029, *t* =−.162, *p* =.87).

To visualize the association between EAA at age 6 and internalizing symptoms from ages 6 to 10, we divided participants into three groups based on EAA scores; EAA>1SD *below* the mean (n =23), EAA>1SD *above* the mean (n =28), and EAA ± 1SD around the mean (n =96). Figure 2 shows the standardized CBCL internalizing scores from ages 6 to 10 for these 3 EAA groups. CBCL data was standardized to compare symptom scores across time. Figure 1B furthermore shows a scatterplot of the cross-sectional association between standardized internalizing symptoms at age 6 and EAA at age 6.

**Figure 2.**
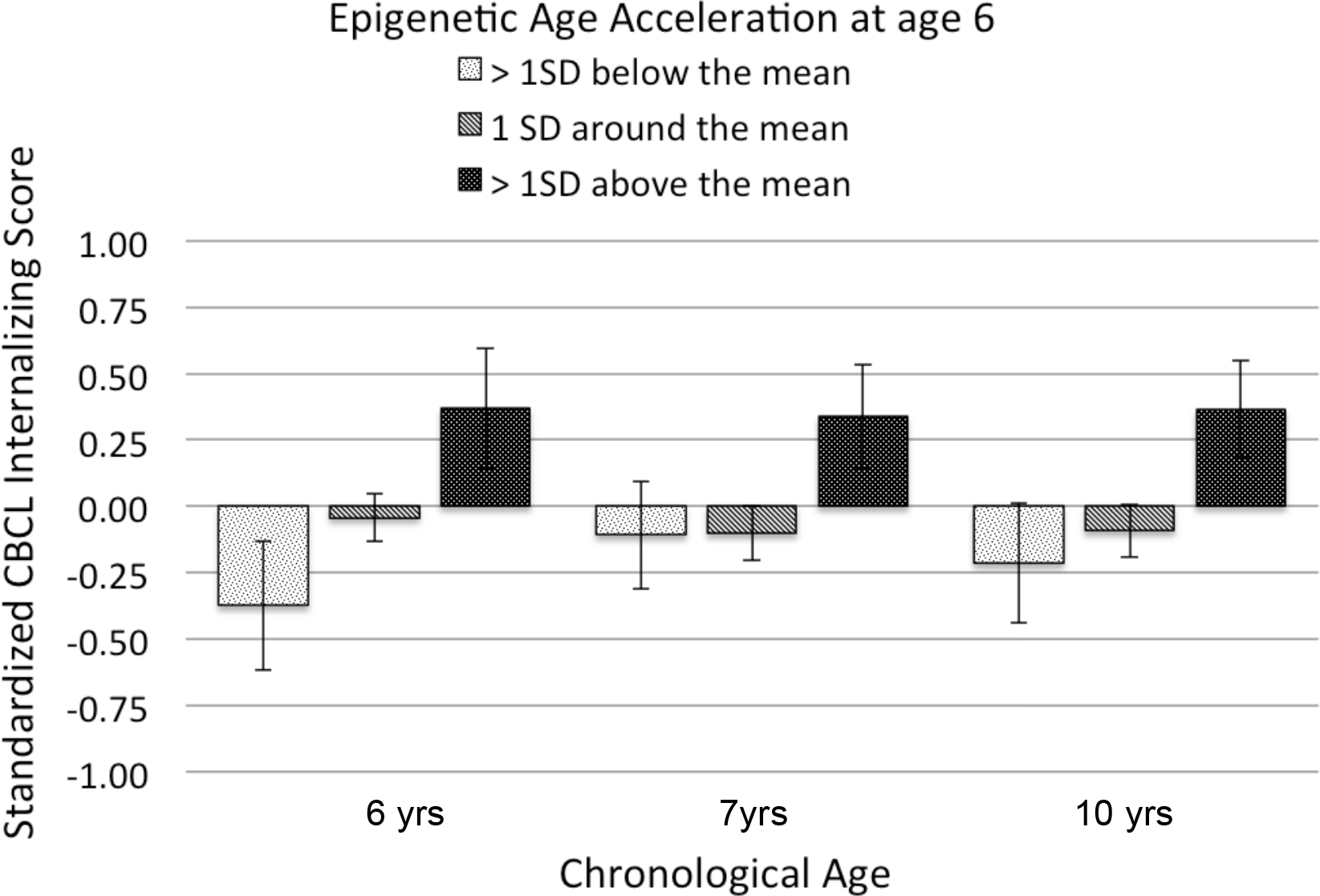
Associations between Epigenetic Age Acceleration (EAA) at age 6, expressed as distance from the mean (EAA>1SD *below* the mean, EAA>1SD *above* the mean, and EAA ± a1SD around the mean), and internalizing symptoms measured with the Child Behavior Checklist (CBCL) over time (6-10 yrs). CBCL values are expressed as standardized scores (*M* =0, *SD* =1) around the mean of the chronological age group. Group differences are indicated by * *p* <.05, ** *p* <.01, yrs = years.

### Externalizing symptoms

CBCL externalizing symptoms did not vary from age 6 to age 10 years (Beta Time =.023, SE =.013, *t* =−1.741, *p* =.083). Maternal STAI scores significantly positively predicted externalizing symptoms over time (beta =.0059, SE =.0026, *t* =2.241, *p* =.026), indicating that maternal anxiety was associated with mothers’ report on child externalizing symptoms. EAA showed a positive, but non-significant association with externalizing symptoms (beta =.062, SE =.033, *t* =1.882, *p* =.062), explaining .8% of variance between individuals. The interaction of EAA with time was not significant either (*p* =.72), see Supplementary TableS8.

In line with the CBCL data, EAA showed a positive, but non-significant, association with SDQ-assessed externalizing scores from age 8 to 10 (beta =.0576, SD =.0296, *t* =1.950, *p* =.053), explaining 2.3% of variance between individuals. The interaction of EAA with time was also not significant (beta =.0446, SD =.024, *t* =1.83, *p* =.069).

## Discussion

We found that advanced biological aging at age 6, assessed by EAA, was significantly associated with internalizing symptoms in children from the age of 2.5 to 10 years. EAA at age 6 was predicted by internalizing symptoms as early as age 2.5, and was longitudinally associated with internalizing symptoms up to the age of 10. Furthermore, EAA and early childhood internalizing symptoms were partly independent predictors of later childhood internalizing symptoms, demonstrating the potential of EAA as an independent biological risk index of childhood psychiatric symptoms. Results were confirmed with a second measure of internalizing symptoms at the ages of 8 and 10, and mothers’ own anxiety and depression scores did not confound these results. The association between EAA and externalizing symptoms in a similar direction did not reach significance in the analyses.

This is one of the first studies to examine associations between epigenetic aging and longitudinally developing psychiatric symptoms in a pediatric sample. A recent study in adolescent girls shows similar associations between EAA and internalizing symptoms (Suarez, Lahti, Czamara, Lahti-Pulkkinen, Girchenko, et al., 2018). Our findings are also in line with studies showing associations between advanced physical maturation and depressive symptoms in childhood (Copeland et al., 2019; Ellis et al., 2019; Lewis et al., 2018). Suarez, Lahti, Czamara, Lahti-Pulkkinen, Knight, et al. (2018) furthermore showed that a lower epigenetic gestational age (GA), which is found to predict worse future health outcomes (Arcangeli, Thilaganathan, Hooper, Khan, & Bhide, 2012), was associated with more child psychiatric problems over time. They calculated epigenetic GA from cord blood with a GA specific epigenetic clock (Knight et al., 2016). Cord blood was not available within the current cohort and as such we cannot determine the relation between epigenetic GA and the Horvath clock, although this would be of interest for future studies.

Future studies should focus on possible underlying mechanisms of the association between EAA and internalizing, and possibly externalizing, symptoms. E.g., early life stress may be associated with the development of psychiatric symptoms (Blair, Glynn, Sandman, & Davis, 2011) as well as accelerated epigenetic aging (Simpkin et al., 2016). Furthermore, internalizing symptoms at age 2.5 predicted EAA at age 6. It may be that children at risk for internalizing symptoms (due to genetic and/or environmental risk factors) experience adverse life events as more stressful than children without these problems. This increased stress exposure may impact biological aging (Zannas et al., 2015). However, as we assessed EAA only at age 6, we cannot exclude the possibility that EAA was already evident at age 2.5 or even earlier. Studies including prospective biosampling from birth with repeated measures of both EAA and childhood psychiatric symptoms over time, or examining interventions aimed at reducing psychopathology or EAA will be important for gaining insight into causality.

The finding that children who exhibit heightened internalizing symptoms also show signs of accelerated biological aging is also of interest given the association between psychiatric disorders and increased risk of aging-related diseases in adulthood (Bennett & Thomas, 2014; Penninx, 2017) and the growing literature associating EAA with early mortality (Chen et al., 2016; Marioni et al., 2015; Perna et al., 2016). Our findings, if replicated, suggest that the burden of internalizing symptoms, even early in childhood, may influence the pace of aging of biological systems, and raise interesting questions about how to reverse such effects (Steve Horvath & Raj, 2018). These findings stress the importance of identifying and treating these symptoms and associated risk factors as early in life as possible.

It is important to note that the current study only examined EAA at the age of 6. It is unknown whether EAA at this age is still associated with EAA and internalizing symptoms in adulthood, and whether it is predictive of physical health symptoms. Longitudinal assessment of psychiatric symptoms, health and EAA are needed to disentangle possible underlying mechanisms. Furthermore, it will be important to integrate other measures of DNA methylation based or cellular aging (Belsky et al., 2017). For example, shortened leukocyte telomere length at age 38 was associated with the persistence of internalizing disorders from ages 11 to 38 years (Shalev et al., 2014).

Epigenetic age estimates in the current study were significantly lower than participants’ chronological age, suggesting a slower pace of epigenetic age in buccal tissue, a feature also observed in neural tissue (Horvath et al., 2015), although this may have been caused by the use of the EPIC array or the normalization measures (McEwen et al., 2018). Future cross-tissue analyses of EAA should determine if buccal tissue is a better proxy for brain-based phenotypes than tissues of a non-ectodermal origin e.g., blood or saliva. While the Horvath clock shows very high accuracy in chronological age prediction (Chen et al., 2016; Horvath, 2013), epigenetic age did not correlate with chronological age in the current sample. This is likely due to the limited age range of the participants during DNA collection, ranging from 5.87 to 6.85 years, but could also be due to the young sample. Previous studies that have used the Horvath-based epigenetic clock in young children also show low correlations with chronological age, but still find associations with developmental outcomes (Simpkin et al., 2016, 2017). This indicates that despite the limited correlations with chronological age in young children, epigenetic aging may still reflect underlying biological aging processes that are informative for child health. The development of specific epigenetic age estimators for pediatric samples will allow for even more precise analyses of biological aging and child development.

Strengths of this study are the prospective longitudinal design with multiple assessments of psychiatric symptoms during childhood. Nonetheless, the sample was relatively small and thereby under-powered to detect small effect sizes, which may explain the non-significant associations with externalizing symptoms and the genetic predictors of EAA. Similarly, sample size precluded stratified analyses based on sex, which may be important (Suarez, Lahti, Czamara, Lahti-Pulkkinen, Knight, et al., 2018). Furthermore, as this is one of the first reports of an association between internalizing symptoms and EAA in childhood it is possible that our relatively large effect sizes represent a ‘winners curse’ and highlight the need for replication in independent and larger samples. Also of note is that maternal report of childhood symptoms was used, which raises concerns about reporter bias, i.e., anxious mothers may view their children as more anxious. However, adjusting our models for maternal anxiety and depression symptoms at each of the time points that child symptoms were assessed did not change our findings. In an optimal design though, objective observer reports or child self-report should be used to assess child behavior over time to disentangle maternal influences. Lastly, our sample includes children with low levels of psychiatric symptoms, coming from relatively highly educated families from a small area in the Netherlands.

## Conclusion

We report a novel association between child EAA and internalizing symptoms across childhood. Future work should determine the importance of this association for long-term (mental) health outcomes. A next important question is whether addressing internalizing symptoms in children, or the stressful environments they experience, will stop the progress of, or even reduce accelerated epigenetic aging. Interestingly, Brody, Yu, Chen, Beach, & Miller (2016) showed that a family-centered prevention program ameliorated the longitudinal association between risky family processes and epigenetic aging. It will also be interesting to examine whether lifestyle programs, including exercise (Quach et al., 2017), might affect EAA and psychiatric symptoms in at-risk children. As EAA has been shown to predict all-cause mortality (Chen et al., 2016), early indicators and prevention of heightened EAA are an important area of further study.

## Acknowledgements

This research was supported by the Netherlands Organization for Scientific Research VIDI grant 575-25-009 (CdeW), a Jacobs Foundation Advanced Research Fellowship (CdeW), and a Sara van Dam Project Grant of the Royal Netherlands Academy of Arts and Sciences (RB). IS was supported by the Mark T. Greenberg Early Career Professorship. KOD is a CIFAR Azrieli Global Scholar in the Child and Brain Development Program. Additional support came from the National Institutes of Health, Eunice Kennedy Shriver National Institute of Child Health and Human Development under Award Number P50HD089922, and the Canada First Research Excellence Fund: Healthy Brains for Healthy Lives.

