## Supplemental information and Tables S1-8 for "Internalizing Symptoms Associate with the Pace of Epigenetic Aging in Childhood"

Content Page

| Content | Page |
|---|---|
| Detailed description of the sample and procedures | 2 |
| TableS1 | 5 |
| TableS2 | 6 |
| TableS3 | 9 |
| TableS4 | 11 |
| TableS5 | 12 |
| TableS6 | 13 |
| TableS7 | 14 |
| TableS8 | 16 |
| Additional references | 17 |

Detailed description of the sample and procedures

The longitudinal study started in 2006 and examines how the environment interacts with individual characteristics to influence development. It originally included 220 mothers in late pregnancy. Only uncomplicated, singleton term pregnancies with 5-min Apgar scores of ≥7 were included in the longitudinal study, leading to a final study sample of 193 mothers and children. At age 6 DNA was sampled in 148 children (69 girls, 79 boys). Of this group, internalizing and externalizing problem symptoms were measured at ages 2.5 years (CBCL n = 145), 4 years (CBCL n = 139), 6 years (CBCL n = 143), 7 years (CBCL n = 143), 8 years (SDQ n = 145) and 10 years (CBCL n = 133, SDQ n = 137). For 116 children data of all measurement time points were available. These children did not differ from those with missing data on demographic, EAA or behavioral measures (all *p*s >.05).

The CBCL 1.5-5 consists of 99 items. The internalizing dimension of this version consists of the scales measuring anxious/depressed, emotionally reactive, somatic, and withdrawn behaviors, and the externalizing dimension consists of the scales measuring attention problems and aggressive behaviors. Continuous scales were calculated, and added to create the two dimension scores. The CBCL 4-18 consists of 112 items. The internalizing dimension of this version consists of the scales measuring anxious/depressed, somatic, and withdrawn behaviors, and the externalizing dimension consists of the scales measuring delinquent and aggressive behaviors. Continuous scales were calculated, and added to create the two dimension scores. The SDQ consists of 25 items. The internalizing dimension consists of the scales measuring emotional symptoms and peer relationship problems, and the externalizing dimension consists of the scales measuring conduct problems and hyperactivity/inattention. Again, scales were added to calculate the two dimension scores. No sex differences were found on the CBCL scales at any age (all *p*s >.07). At age 10 only, girls exhibited higher levels of SDQ-reported internalizing problems than boys ($F[1, 135] = 4.03$, $p = .04$).

Genomic DNA was extracted from buccal epithelial cells with the QIAamp DNA Mini Kit (Qiagen, Germany) and quantified using a Nanodrop 2000 spectrophotometer (Thermo Fisher Scientific). Next, 750ng of gDNA was bisulphite converted using the Zymo EZ dna methylation kit (Zymo Research, Irvine, CA, USA), and genome-wide DNA methylation described using the Infinium EPIC array (850K array; Illumina, San Diego CA, USA) according to manufacturers guidelines.

Genotyping was performed on the same buccal samples using the Infinium Global Screening Array (Illumina, Inc.). SNPs with missing rates over 5% and with p-values less than 1e-20 on the Hardy-Weinberg Equilibrium exact test were excluded. Two participants were excluded as they had missing SNP rates higher than 5%. The Sanger Imputation Service with the Haplotype Reference Consortium (release 1.1) was used to impute missing genotypes (McCarthy et al., 2016). Only SNPs with high imputation accuracy (INFO Score ≥ .80) were considered in the genetic analyses.

Raw DNA methylation beta values derived from buccal cells at age 6 were uploaded to https://dnamage.genetics.ucla.edu, using the 'normalize data' option, together with a phenotype file including precise chronological age, to calculate the DNA methylation (DNAm) based age and EAA. DNAm age estimates using normal-exponential out-of-band (NOOB) background corrections gave almost similar results.

Potential genetic risk factors for EAA were considered. Two recent GWAS provide evidence of a genetic contribution to EAA in the brain (Lu et al., 2016) and in blood (Lu et al., 2018). To date no GWAS of EAA in buccal tissue is available. Therefore, SNPs that were genome-wide significantly associated with EAA in brain or blood were examined as possible covariates in the analyses predicting EAA at age 6 (5 brain-based SNPs: rs30986, rs1128156, rs2054847, rs6723868, rs7305937; 7 blood-based SNPs: rs11706810, rs2736099, rs143093668, rs6915893, rs73397619, rs78781855, rs1005277). However, none of these SNPs contributed to EAA (see TableS2) and were therefore not included in further analyses.

S1. Spearman rank correlations between internalizing and externalizing symptom scales over time.

| Child Outcomes | | CBCL INT 2.5y | CBCL INT 4y | CBCL INT 6y | CBCL INT 7y | CBCL INT 10y | SDQ INT 8y | SDQ INT 10y |
|---|---|---|---|---|---|---|---|---|
| CBCL EXT 2.5y | Rho | .547** | .375** | .259** | .226** | .276** | .187* | .205* |
| | *p*-value | .000 | .000 | .002 | .007 | .001 | .026 | .018 |
| | N | 143 | 135 | 138 | 139 | 130 | 141 | 133 |
| CBCL EXT 4y | Rho | .376** | .598** | .288** | .185* | .377** | .158 | .259** |
| | *p*-value | .000 | .000 | .001 | .032 | .000 | .068 | .003 |
| | N | 136 | 138 | 135 | 134 | 124 | 135 | 128 |
| CBCL EXT 6y | Rho | .369** | .370** | .462** | .305** | .441** | .193* | .256** |
| | *p*-value | .000 | .000 | .000 | .000 | .000 | .022 | .003 |
| | N | 139 | 135 | 142 | 139 | 129 | 140 | 134 |
| CBCL EXT 7y | Rho | .282** | .348** | .342** | .428** | .434** | .234** | .228** |
| | *p*-value | .001 | .000 | .000 | .000 | .000 | .005 | .008 |
| | N | 140 | 134 | 139 | 142 | 130 | 141 | 134 |
| CBCL EXT 10y | Rho | .266** | .323** | .207* | .215* | .452** | .152 | .186* |
| | *p*-value | .002 | .000 | .018 | .014 | .000 | .082 | .036 |
| | N | 130 | 124 | 129 | 130 | 132 | 132 | 127 |
| SDQ EXT 8y | Rho | .189* | .140 | .061 | .178* | .315** | .207* | .202* |
| | *p*-value | .024 | .105 | .475 | .035 | .000 | .013 | .018 |
| | N | 142 | 135 | 140 | 141 | 132 | 144 | 136 |
| SDQ EXT 10y | Rho | .205* | .188* | .078 | .155 | .300** | .121 | .244** |
| | *p*-value | .017 | .033 | .370 | .073 | .001 | .159 | .004 |
| | N | 134 | 128 | 134 | 134 | 127 | 136 | 136 |

Note: y = years. EAA = Epigenetic Age Acceleration, CBCL = Child Behavior Checklist, SDQ = Strengths and Difficulties Questionnaire, INT = internalizing, EXT = externalizing. * $p<.05$, ** $p<.01$.

S2. Spearman rank correlations between EAA, CBCL, SDQ, and demographic, technical and genetic covariates (*continues on next page*)

| Child Outcomes | | EAA | CBCL INT 2.5y | CBCL EXT 2.5y | CBCL INT 4y | CBCL EXT 4y | CBCL INT 6y | CBCL EXT 6y | CBCL INT 7y | CBCL EXT 7y | CBCL INT 10y | CBCL EXT 10y | SDQ INT 8y | SDQ EXT 8y | SDQ INT 10y | SDQ EXT 10y |
|---|---|---|---|---|---|---|---|---|---|---|---|---|---|---|---|---|
| *Covariates* | | | | | | | | | | | | | | | | |
| | Rho | -.044 | .154 | .069 | .068 | .109 | .009 | .056 | -.078 | .068 | .046 | .036 | .058 | .065 | .173* | .049 |
| Gender | *p*-value | .599 | .066 | .411 | .429 | .203 | .917 | .507 | .357 | .424 | .603 | .683 | .489 | .441 | .044 | .571 |
| | N | 147 | 144 | 143 | 138 | 138 | 142 | 142 | 142 | 142 | 132 | 132 | 144 | 144 | 136 | 136 |
| | Rho | .057 | .042 | .063 | -.067 | -.098 | -.029 | -.063 | .023 | -.018 | -.032 | -.071 | .018 | .015 | -.092 | -.067 |
| Maternal Education | *p*-value | .500 | .626 | .459 | .439 | .260 | .737 | .463 | .784 | .837 | .721 | .428 | .837 | .857 | .293 | .448 |
| | N | 143 | 140 | 139 | 134 | 134 | 138 | 138 | 139 | 139 | 128 | 128 | 140 | 140 | 132 | 132 |
| | Rho | .022 | -.050 | -.064 | .108 | .030 | -.001 | -.097 | .060 | -.093 | .092 | .015 | **.198*** | -.068 | .109 | -.113 |
| Batch | *p*-value | .792 | .554 | .449 | .206 | .723 | .988 | .249 | .475 | .269 | .296 | .866 | **.017** | .415 | .207 | .191 |
| | N | 147 | 144 | 143 | 138 | 138 | 142 | 142 | 142 | 142 | 132 | 132 | **144** | 144 | 136 | 136 |
| | Rho | .101 | -.001 | -.029 | .156 | .032 | .117 | -.083 | **.168*** | -.075 | .105 | -.028 | **.193*** | -.123 | .186* | -.142 |
| Array plate | *p*-value | .225 | .989 | .727 | .067 | .706 | .164 | .325 | **.045** | .374 | .229 | .748 | **.021** | .141 | .030 | .100 |
| | N | 147 | 144 | 143 | 138 | 138 | 142 | 142 | **142** | 142 | 132 | 132 | **144** | 144 | 136 | 136 |
| | Rho | **-.191*** | .100 | .020 | -.080 | -.076 | .026 | -.016 | -.033 | .040 | -.045 | .031 | .062 | -.063 | .061 | -.034 |
| Array position | *p*-value | **.020** | .234 | .813 | .351 | .376 | .755 | .848 | .694 | .633 | .608 | .724 | .462 | .454 | .484 | .695 |
| | N | **147** | 144 | 143 | 138 | 138 | 142 | 142 | 142 | 142 | 132 | 132 | 144 | 144 | 136 | 136 |
| Buccal Cell count | Rho | .036 | -.003 | .000 | .021 | **.178*** | .045 | .151 | .039 | .120 | .068 | .016 | .038 | -.074 | -.075 | .011 |
| | *p*-value | .669 | .973 | .996 | .811 | **.037** | .599 | .072 | .648 | .156 | .442 | .856 | .653 | .379 | .387 | .898 |
| | N | 147 | 144 | 143 | 138 | **138** | 142 | 142 | 142 | 142 | 132 | 132 | 144 | 144 | 136 | 136 |
| CD34 Cell count | Rho | -.069 | -.038 | -.017 | -.018 | -.147 | -.110 | **-.182*** | -.101 | -.156 | -.130 | -.070 | -.096 | .036 | .015 | -.007 |
| | *p*-value | .405 | .655 | .844 | .834 | .085 | .192 | **.030** | .232 | .064 | .136 | .427 | .254 | .668 | .862 | .935 |
| | N | 147 | 144 | 143 | 138 | 138 | 142 | **142** | 142 | 142 | 132 | 132 | 144 | 144 | 136 | 136 |
| | Rho | .014 | -.098 | -.019 | -.123 | .009 | **-.178*** | -.007 | -.038 | -.021 | -.030 | .024 | .002 | .143 | .030 | .070 |
| PC1 | *p*-value | .867 | .246 | .823 | .155 | .913 | **.035** | .934 | .652 | .809 | .739 | .785 | .977 | .090 | .732 | .421 |
| | N | 145 | 142 | 141 | 136 | 136 | **140** | 140 | 141 | 141 | 130 | 130 | 142 | 142 | 134 | 134 |
| | Rho | -.096 | -.118 | -.079 | -.014 | -.076 | -.056 | -.119 | .041 | -.025 | .022 | .018 | .094 | .083 | .024 | .009 |
| PC2 | *p*-value | .251 | .160 | .352 | .869 | .382 | .510 | .162 | .628 | .772 | .800 | .838 | .267 | .326 | .786 | .916 |
| | N | 145 | 142 | 141 | 136 | 136 | 140 | 140 | 141 | 141 | 130 | 130 | 142 | 142 | 134 | 134 |

| | | | | | | | | | | | | | | | | |
|---|---|---|---|---|---|---|---|---|---|---|---|---|---|---|---|---|
| *EAA related brain-based SNPs:* | | | | | | | | | | | | | | | | |
| rs30986 | Rho | -.088 | .062 | -.013 | .046 | .073 | .047 | .017 | -.036 | -.045 | .094 | .022 | .000 | -.042 | .088 | -.031 |
| | *p*-value | .333 | .505 | .891 | .622 | .438 | .613 | .855 | .700 | .629 | .329 | .817 | .999 | .649 | .355 | .746 |
| | N | 122 | 119 | 119 | 115 | 115 | 119 | 119 | 118 | 118 | 110 | 110 | 119 | 119 | 112 | 112 |
| rs1128156 | Rho | .018 | -.027 | -.052 | -.016 | -.013 | .097 | -.023 | -.022 | -.048 | .016 | -.096 | -.006 | -.059 | **.171*** | .009 |
| | *p*-value | .831 | .752 | .540 | .857 | .884 | .253 | .784 | .793 | .573 | .853 | .278 | .943 | .486 | **.048** | .915 |
| | N | 145 | 142 | 141 | 136 | 136 | 140 | 140 | 141 | 141 | 130 | 130 | 142 | 142 | **134** | 134 |
| rs2054847 | Rho | -.004 | -.003 | -.060 | .024 | -.010 | .021 | -.042 | -.036 | -.066 | .026 | -.107 | -.003 | -.062 | .154 | -.039 |
| | *p*-value | .963 | .971 | .476 | .781 | .904 | .805 | .626 | .668 | .434 | .767 | .227 | .974 | .464 | .076 | .658 |
| | N | 145 | 142 | 141 | 136 | 136 | 140 | 140 | 141 | 141 | 130 | 130 | 142 | 142 | 134 | 134 |
| rs6723868 | Rho | .002 | .043 | -.105 | .002 | -.084 | -.086 | -.082 | .003 | -.069 | .056 | .074 | .051 | -.049 | .064 | -.010 |
| | *p*-value | .980 | .626 | .233 | .979 | .351 | .330 | .358 | .970 | .431 | .542 | .420 | .563 | .577 | .478 | .909 |
| | N | 134 | 131 | 130 | 126 | 126 | 129 | 129 | 131 | 131 | 120 | 120 | 131 | 131 | 124 | 124 |
| rs7305937 | Rho | .154 | -.007 | -.077 | -.004 | -.014 | .101 | .070 | .036 | .129 | .069 | .102 | .077 | .044 | -.008 | .081 |
| | *p*-value | .064 | .931 | .366 | .965 | .876 | .235 | .412 | .669 | .129 | .439 | .249 | .362 | .603 | .931 | .353 |
| | N | 145 | 142 | 141 | 136 | 136 | 140 | 140 | 141 | 141 | 130 | 130 | 142 | 142 | 134 | 134 |

*(TableS2 continued)*

| Child Outcomes | | EAA | CBCL INT 2.5y | CBCL EXT 2.5y | CBCL INT 4y | CBCL EXT 4y | CBCL INT 6y | CBCL EXT 6y | CBCL INT 7y | CBCL EXT 7y | CBCL INT 10y | CBCL EXT 10y | SDQ INT 8y | SDQ EXT 8y | SDQ INT 10y | SDQ EXT 10y |
|---|---|---|---|---|---|---|---|---|---|---|---|---|---|---|---|---|
| *EAA related blood-based SNPs:* | | | | | | | | | | | | | | | | |
| rs11706810 | Rho | -.065 | -.057 | -.127 | -.007 | .018 | .067 | .073 | -.040 | -.034 | .047 | .067 | **-.206*** | -.050 | -.022 | .018 |
| | *p*-value | .440 | .502 | .134 | .935 | .832 | .434 | .394 | .634 | .691 | .592 | .450 | **.014** | .556 | .805 | .840 |
| | N | 144 | 141 | 140 | 136 | 136 | 140 | 140 | 141 | 141 | 130 | 130 | **141** | 141 | 134 | 134 |
| rs2736099 | Rho | .031 | -.009 | .080 | .061 | .114 | -.154 | .030 | -.114 | -.050 | -.113 | -.030 | .013 | .015 | .020 | .076 |
| | *p*-value | .725 | .916 | .367 | .497 | .206 | .081 | .734 | .195 | .569 | .219 | .743 | .881 | .863 | .826 | .404 |
| | N | 134 | 131 | 130 | 125 | 125 | 129 | 129 | 130 | 130 | 121 | 121 | 131 | 131 | 123 | 123 |
| rs143093668 | Rho | -.034 | .000 | -.041 | .082 | .142 | .088 | .087 | .009 | .004 | .146 | .123 | -.020 | -.118 | -.047 | -.169 |
| | *p*-value | .683 | .996 | .634 | .344 | .102 | .305 | .310 | .914 | .964 | .098 | .164 | .817 | .164 | .594 | .053 |

| | | | | | | | | | | | | | | | | |
|---|---|---|---|---|---|---|---|---|---|---|---|---|---|---|---|---|
| | N | 143 | 140 | 139 | 134 | 134 | 138 | 138 | 139 | 139 | 129 | 129 | 140 | 140 | 132 | 132 |
| | Rho | -.054 | .057 | .028 | .101 | .103 | .067 | -.011 | -.031 | -.010 | -.106 | .021 | -.091 | .055 | .135 | **.193*** |
| rs6915893 | *p*-value | .515 | .503 | .739 | .241 | .234 | .433 | .898 | .712 | .903 | .231 | .815 | .282 | .516 | .121 | **.025** |
| | N | 145 | 142 | 141 | 136 | 136 | 140 | 140 | 141 | 141 | 130 | 130 | 142 | 142 | 134 | **134** |
| | Rho | -.003 | .011 | .069 | -.101 | .011 | .108 | .071 | .004 | .023 | .013 | -.022 | -.093 | .124 | -.083 | .116 |
| rs73397619 | *p*-value | .967 | .892 | .413 | .240 | .896 | .202 | .404 | .966 | .789 | .879 | .808 | .271 | .143 | .340 | .181 |
| | N | 145 | 142 | 141 | 136 | 136 | 140 | 140 | 141 | 141 | 130 | 130 | 142 | 142 | 134 | 134 |
| | Rho | .153 | .164 | .148 | .169 | .052 | .071 | .045 | .066 | .046 | .030 | .049 | -.061 | .033 | -.162 | -.041 |
| rs78781855 | *p*-value | .070 | .053 | .083 | .052 | .555 | .413 | .606 | .442 | .594 | .734 | .585 | .478 | .701 | .064 | .638 |
| | N | 141 | 139 | 138 | 132 | 132 | 136 | 136 | 138 | 138 | 128 | 128 | 139 | 139 | 131 | 131 |
| | Rho | .034 | -.022 | -.096 | .019 | -.124 | .055 | .082 | .075 | .013 | -.043 | .040 | .159 | -.137 | **.200*** | -.024 |
| rs1005277 | *p*-value | .690 | .801 | .261 | .828 | .153 | .524 | .339 | .377 | .880 | .626 | .657 | .061 | .108 | **.021** | .783 |
| | N | 143 | 140 | 139 | 134 | 134 | 138 | 138 | 139 | 139 | 128 | 128 | 140 | 140 | **132** | 132 |

Note: [a] 0 female / 1 = male, y = years, EAA = Epigenetic Age Acceleration, CBCL = Child Behavior Checklist, SDQ = Strengths and Difficulties Questionnaire, INT = internalizing, EXT = externalizing, STAI = State Anxiety scores (State-Trait Anxiety Inventory), EPDS = State Depression scores (Edinburgh Postnatal Depression Scale), * $p<.05$, ** $p<.01$, significant findings are printed in bold. Only few significant associations between child outcomes and covariates were found.

S3. Spearman rank correlations between internalizing and externalizing symptom scales and maternal STAI and EPDS scores at all ages

| Child Outcomes | | EAA | CBCL INT 2.5y | CBCL EXT 2.5y | CBCL INT 4y | CBCL EXT 4y | CBCL INT 6y | CBCL EXT 6y | CBCL INT 7y | CBCL EXT 7y | CBCL INT 10y | CBCL EXT 10y | SDQ INT 8y | SDQ EXT 10y |
|---|---|---|---|---|---|---|---|---|---|---|---|---|---|---|
| Maternal Mood | | | | | | | | | | | | | | |
| STAI 2.5y | Rho | -.066 | .173* | .069 | .163 | .200* | .222** | .185* | .252** | .153 | .265** | .210* | .368** | .183* |
| | *p*-value | .434 | .038 | .413 | .058 | .019 | .009 | .030 | .003 | .072 | .002 | .016 | .000 | .029 |
| | N | 144 | 144 | 143 | 136 | 136 | 139 | 139 | 140 | 140 | 130 | 130 | 142 | 142 |
| EPDS 2.5y | Rho | -.030 | .337** | .195* | .296** | .220* | .119 | .151 | .228** | .183* | .279** | .222* | .244** | .098 |
| | *p*-value | .719 | .000 | .020 | .001 | .010 | .165 | .076 | .007 | .031 | .001 | .011 | .004 | .245 |
| | N | 143 | 143 | 142 | 135 | 135 | 138 | 138 | 139 | 139 | 130 | 130 | 141 | 141 |
| STAI 4y | Rho | -.032 | .056 | .023 | .147 | .129 | .150 | .119 | .233** | .122 | .182* | .110 | .215* | .022 |
| | *p*-value | .709 | .512 | .787 | .090 | .137 | .082 | .169 | .007 | .159 | .042 | .223 | .012 | .798 |
| | N | 139 | 137 | 136 | 135 | 135 | 136 | 136 | 135 | 135 | 125 | 125 | 136 | 136 |
| EPDS 4y | Rho | -.052 | .206* | -.029 | .207* | .161 | .121 | .041 | .095 | .117 | .183* | .041 | .033 | -.024 |
| | *p*-value | .544 | .015 | .738 | .015 | .061 | .156 | .633 | .269 | .174 | .040 | .646 | .700 | .782 |
| | N | 141 | 139 | 138 | 137 | 137 | 138 | 138 | 137 | 137 | 127 | 127 | 138 | 138 |
| STAI 6y | Rho | -.092 | .102 | .042 | .111 | .097 | .250** | .145 | .180* | .058 | .160 | .047 | .226** | .012 |
| | *p*-value | .281 | .234 | .627 | .202 | .263 | .003 | .087 | .035 | .501 | .071 | .600 | .008 | .886 |
| | N | 141 | 138 | 138 | 134 | 134 | 141 | 141 | 138 | 138 | 129 | 129 | 139 | 139 |
| EPDS 6y | Rho | -.099 | .185* | .109 | .075 | .175* | .090 | .093 | .122 | .084 | .148 | .071 | .160 | .116 |
| | *p*-value | .244 | .030 | .206 | .391 | .044 | .289 | .274 | .155 | .329 | .096 | .424 | .062 | .176 |
| | N | 140 | 137 | 137 | 133 | 133 | 140 | 140 | 137 | 137 | 128 | 128 | 138 | 138 |
| STAI 8y | Rho | .056 | .144 | .044 | .143 | .125 | .187* | .178* | .297** | .155 | .261** | .134 | .374** | .119 |
| | *p*-value | .506 | .089 | .609 | .100 | .151 | .027 | .036 | .000 | .067 | .003 | .126 | .000 | .158 |
| | N | 143 | 141 | 140 | 134 | 134 | 139 | 139 | 140 | 140 | 131 | 131 | 143 | 143 |
| EPDS 8y | Rho | .060 | .205* | .143 | .158 | .165 | .124 | .172* | .233** | .154 | .252** | .211* | .330** | .156 |
| | *p*-value | .480 | .015 | .095 | .070 | .059 | .149 | .044 | .006 | .071 | .004 | .016 | .000 | .065 |
| | N | 141 | 139 | 138 | 132 | 132 | 137 | 137 | 138 | 138 | 129 | 129 | 141 | 141 |
| STAI 10y | Rho | -.004 | .153 | .124 | .138 | .226* | .273** | .128 | .282** | .158 | .303** | .185* | .400** | .119 |
| | *p*-value | .962 | .082 | .157 | .123 | .011 | .002 | .147 | .001 | .072 | .000 | .034 | .000 | .171 |
| | N | 133 | 131 | 131 | 125 | 125 | 130 | 130 | 131 | 131 | 132 | 132 | 133 | 133 |

| | | | | | | | | | | | | | | |
|---|---|---|---|---|---|---|---|---|---|---|---|---|---|---|
| EPDS | Rho | -.048 | .234** | .231** | .183* | .214* | .189* | .091 | .219* | .096 | .236** | .133 | .311** | .080 |
| 10y | *p*-value | .582 | .007 | .008 | .041 | .017 | .031 | .304 | .012 | .277 | .006 | .127 | .000 | .360 |
| | N | 133 | 131 | 131 | 125 | 125 | 130 | 130 | 131 | 131 | 132 | 132 | 133 | 133 |

Note: y = years, EAA = Epigenetic Age Acceleration, CBCL = Child Behavior Checklist, SDQ = Strengths and Difficulties Questionnaire, INT = internalizing, EXT = externalizing, STAI = State Anxiety scores (State-Trait Anxiety Inventory), EPDS = State Depression scores (Edinburgh Postnatal Depression Scale), * $p<.05$, ** $p<.01$.

S4. Regression models with CBCL internalizing symptoms at ages 2.5 and 4 predicting EAA at age 6.

| Outcome: | EAA at age 6 | | | | | | | | | | | | | | | |
|---|---|---|---|---|---|---|---|---|---|---|---|---|---|---|---|---|
| | Model1 | | | | Model 2 | | | | Model 3 | | | | Model 4 | | | |
| | B | SE | Beta | *p* | B | SE | Beta | *p* | B | SE | Beta | *p* | B | SE | Beta | *p* |
| Predictors: | | | | | | | | | | | | | | | | |
| Array position | -.062 | .029 | -.178* | .033 | -.065 | .028 | -.188* | .022 | -.071 | .028 | -208* | .013 | -.080 | .029 | -.233** | .006 |
| CBCL 2.5yrs | - | - | - | - | .524 | .213 | .200* | .015 | - | - | - | - | .562 | .268 | .215* | .038 |
| CBCL 4 yrs | - | - | - | - | - | - | - | - | .455 | .200 | .189* | .024 | .159 | .249 | .066 | .525 |
| $R^2$ | .032* | | | | .072** | | | | .087** | | | | .118*** | | | |
| $\Delta R^2$ | | | | | .040* | | | | 035* | | | | .003 | | | |

Note: EAA = Epigenetic Age Acceleration, CBCL = Child Behavior Checklist, yrs = years, * *p*<.05, ** *p*<.01, *** *p*<.001.

S5. Regression models with CBCL externalizing symptoms at ages 2.5 and 4 predicting EAA at age 6.

| Outcome: | EAA at age 6 | | | | | | | | | | | | | | | |
|---|---|---|---|---|---|---|---|---|---|---|---|---|---|---|---|---|
| | Model1 | | | | Model 2 | | | | Model 3 | | | | Model 4 | | | |
| | B | SE | Beta | *p* | B | SE | Beta | *p* | B | SE | Beta | *p* | B | SE | Beta | *p* |
| Predictors: | | | | | | | | | | | | | | | | |
| Array position | -.063 | .029 | -.180* | .031 | -.064 | .029 | -.183* | .028 | -.073 | .029 | -.214* | .012 | -.082 | .029 | -.238** | .006 |
| CBCL 2.5yrs | - | - | - | - | .397 | .266 | .123 | .137 | - | - | - | - | .647 | .338 | .198 | .058 |
| CBCL 4 yrs | - | - | - | - | - | - | - | - | .285 | .201 | .119 | .158 | <.001 | .253 | <.001 | 1.00 |
| $R^2$ | .033* | | | | .048* | | | | .066** | | | | .092** | | | |
| $\Delta R^2$ | | | | | .015 | | | | .014 | | | | <.001 | | | |

Note: EAA = Epigenetic Age Acceleration, CBCL = Child Behavior Checklist, yrs = years, * *p*<.05, ** *p*<.01, *** *p*<.001.

S6. Mixed model analyses with EAA at age 6 predicting longitudinal CBCL internalizing symptoms from ages 6 to 10.

| Outcome: | Internalizing symptoms from age 6-10 | | | | | | | | | | | |
|---|---|---|---|---|---|---|---|---|---|---|---|---|
| | Model 1 | | | | Model 2 | | | | Model 3 | | | |
| | B | SE | t | *p* | B | SE | t | *p* | B | SE | t | *p* |
| Predictors: | | | | | | | | | | | | |
| Intercept | .3931 | .0645 | 6.093*** | <.001 | .3871 | .0634 | 6.105*** | <.001 | .3860 | .0634 | 6.079*** | <.001 |
| Time | .0482 | .0134 | 3.587*** | <.001 | .0482 | .0134 | 3.595*** | <.001 | .0482 | .0134 | 3.593*** | <.001 |
| STAI | .00726 | .00195 | 3.729*** | <.001 | .00747 | .00192 | 3.891*** | <.001 | .00751 | .00192 | 3.905*** | <.001 |
| PC1 | -.778 | .286 | -2.72** | .007 | -.770 | .276 | -2.788** | .006 | -.7699 | .2763 | -2.786** | .006 |
| EAA | - | - | - | - | .0959 | .0281 | 3.418*** | .001 | .0955 | .0281 | 3.403*** | .001 |
| EAA*Time | - | - | - | - | - | - | - | - | -.00773 | .0172 | -.450 | .653 |
| BIC | 162.19 | | | | 156.23 | | | | 162.31 | | | |
| σ2r (SE) | .0472 (.0041) | | | | .0473 (.0041) | | | | .0474 (.0042) | | | |
| σ2μ (SE) | .0602 (.0094) | | | | .0545 (.0087) | | | | .0544 (.0087) | | | |
| Δσ2μ (%) | | | | | 9.4 (vs model 1) | | | | .2 (vs model 2) | | | |

Note: EAA = Epigenetic Age Acceleration, CBCL = Child Behavior Checklist, STAI = State Anxiety scores (State-Trait Anxiety Inventory), yrs = years, σ2r = Residual Variance, σ2μ = Intercept Variance, Δσ2μ (%) = $((\sigma 2\mu_{mx} - \sigma 2\mu_{mx+1}) / \sigma 2\mu_{mx})$x100, * *p*<.05, ** *p*<.01, *** *p*<.001.

S7. Mixed model analyses with early childhood symptoms and EAA at age 6 predicting longitudinal CBCL internalizing symptoms from ages 6 to 10.

| Outcome: | Internalizing symptoms from age 6-10 | | | | | | | | | | | | | | | |
|---|---|---|---|---|---|---|---|---|---|---|---|---|---|---|---|---|
| | Model1 | | | | Model 2 | | | | Model 3 | | | | Model 4 | | | |
| | B | SE | t | *p* | B | SE | t | *p* | B | SE | t | *p* | B | SE | t | *p* |
| Predictors: | | | | | | | | | | | | | | | | |
| Time | .047794 | .013558 | 3.525*** | .000 | .047797 | .013565 | 3.523*** | .001 | .050645 | .014031 | 3.609*** | .000 | .050729 | .014039 | 3.613*** | .000 |
| STAI | .006327 | .001903 | 3.325*** | .001 | .006555 | .001885 | 3.478*** | .001 | .005636 | .001909 | 2.953** | .003 | .005890 | .001904 | 3.093** | .002 |
| PC1 | -.576579 | .267342 | -2.157* | .033 | -.594923 | .261213 | -2.278* | .024 | -.667498 | .260031 | -2.567* | .011 | -.661228 | .257452 | -2.568* | .011 |
| CBCL 2.5yrs | .114034 | .021922 | 5.202*** | .000 | .101403 | .021849 | 4.641*** | .000 | .072153 | .026100 | 2.765** | .007 | .064555 | .026129 | 2.471* | .015 |
| CBCL 4yrs | - | - | - | - | - | - | - | - | .092305 | .025881 | 3.567*** | .001 | .087469 | .025723 | 3.400*** | .001 |
| EAA | - | - | - | - | .073938 | .026686 | 2.771** | .006 | - | - | - | - | .050425 | .026864 | 1.877 | .063 |
| BIC | 142.41 | | | | 140.31 | | | | 132.01 | | | | 133.93 | | | |
| σ2r (SE) | .0476 (.0042) | | | | .0476 (.0042) | | | | .0485 (.0044) | | | | 0486 (.0044) | | | |
| σ2μ (SE) | .0477 (.0080) | | | | .0445 (.0077) | | | | .04075 (.0074) | | | | .03946 (.0073) | | | |
| Δσ2μ (%) | | | | | 6.7 (vs model 1) | | | | | | | | 3.1 (vs model 3) | | | |

Note: EAA = Epigenetic Age Acceleration, CBCL = Child Behavior Checklist, STAI = State Anxiety scores (State-Trait Anxiety Inventory), yrs = years, σ2r = Residual Variance, = Intercept Variance, Δσ2μ (%) = ((σ2$\mu_{mx}$ - σ2$\mu_{mx+1)/}$ σ2$\mu_{mx)}$*100, * *p*<.05, ** *p*<.01, *** *p*<.001.

S8. Mixed model analyses with EAA at age 6 predicting longitudinal CBCL externalizing symptoms from ages 6 to 10.

| Outcome: | Externalizing symptoms from age 6-10 | | | | | | | | | | | |
|---|---|---|---|---|---|---|---|---|---|---|---|---|
| | Model 1 | | | | Model 2 | | | | Model 3 | | | |
| Predictors | B | SE | t | *p* | B | SE | t | *p* | B | SE | t | *p* |
| STAI | .004889 | .001951 | 2.505* | .013 | .004934 | .001946 | 2.535* | .012 | .004927 | .001946 | 2.531* | .012 |
| EAA | - | - | - | - | .062484 | .033198 | 1.882 | .062 | .062401 | .033177 | 1.881 | .062 |
| Time | - | - | - | - | - | - | - | - | -.020141 | .012650 | -1.592 | .113 |
| EAA*Time | - | - | - | - | - | - | - | - | -.005961 | .016299 | -.366 | .715 |
| BIC | 171.64 | | | | 173.09 | | | | 183.70 | | | |
| σ2r (SE) | .0427 (.0037) | | | | 0427 (.0037) | | | | 0427 (.0037) | | | |
| σ2μ (SE) | .0866 (.0121) | | | | .0849 (.0119) | | | | .0847 (.0119) | | | |
| Δσ2μ (%) | | | | | 2.0 (vs model 1) | | | | 1.4 (vs model 2) | | | |

Note: EAA = Epigenetic Age Acceleration, CBCL = Child Behavior Checklist, STAI = State Anxiety scores (State-Trait Anxiety Inventory), yrs = years, σ2r = Residual Variance, = Intercept Variance, Δσ2μ (%) = $((\sigma 2\mu_{mx} - \sigma 2\mu_{mx+1})/ \sigma 2\mu_{mx})*100$, * $p$<.05, ** $p$<.01, *** $p$<.001.
